# Contrastive learning on protein embeddings enlightens midnight zone

**DOI:** 10.1101/2021.11.14.468528

**Authors:** Michael Heinzinger, Maria Littmann, Ian Sillitoe, Nicola Bordin, Christine Orengo, Burkhard Rost

## Abstract

Experimental structures are leveraged through multiple sequence alignments, or more generally through homology-based inference (HBI), facilitating the transfer of information from a protein with known annotation to a query without any annotation. A recent alternative expands the concept of HBI from sequence-distance lookup to embedding-based annotation transfer (EAT). These embeddings are derived from protein Language Models (pLMs). Here, we introduce using single protein representations from pLMs for contrastive learning. This learning procedure creates a new set of embeddings that optimizes constraints captured by hierarchical classifications of protein 3D structures defined by the CATH resource. The approach, dubbed *ProtTucker*, has an improved ability to recognize distant homologous relationships than more traditional techniques such as threading or fold recognition. Thus, these embeddings have allowed sequence comparison to step into the “midnight zone” of protein similarity, i.e., the region in which distantly related sequences have a seemingly random pairwise sequence similarity. The novelty of this work is in the particular combination of tools and sampling techniques that ascertained good performance comparable or better to existing state-of-the-art sequence comparison methods. Additionally, since this method does not need to generate alignments it is also orders of magnitudes faster. The code is available at https://github.com/Rostlab/EAT.

## Introduction

### Phase-transition from daylight through twilight into midnight zone

Protein sequence determines structure which determines function. This simple chain underlies the success of grouping proteins into families from sequence (3,19-21). Information from experimental high-resolution three-dimensional (3D) structures expands the perspective from families to super-families (1,22) that often reveal evolutionary and functional connections not recognizable from sequence alone (23,24). Thus, 3D information helps us to penetrate through the twilight zone of sequence alignments (25,26) into the midnight zone of distant evolutionary relationships (27).

The transition from daylight, through twilight and into the midnight zone is characterized by a phase-transition, i.e., a sigmoid function describing an order of magnitude increase in recall (relations identified) at the expense of a decrease in precision (relations identified correctly) over a narrow range of sequence similarity. Measuring sequence similarity by the HSSP-value (<u>HVAL</u>) (26,28) for the *daylight zone* at HVAL>5 (>25% PIDE - pairwise sequence identity over >250 aligned residues) over 90% of all protein pairs have similar 3D structures, while at the beginning of the *midnight zone* for HVAL<-5 (<15% PIDE for >250 aligned residues), over 90% have different 3D structures. Thus, the transition from daylight to midnight zone is described by a phase-transition in which over about ten percentage points in PIDE precision drops from 90% to 10%, i.e., from almost *all correct* to almost *all incorrect* within ±5 points PIDE. The particular point at which the twilight zone begins and how extreme the transition is, depends on the phenotype: steeper at lower PIDE for structure (26) and flatter at higher PIDE for function (29,30).

If two proteins have highly similar structures, it is still possible for their sequences to be found in this *midnight zone*, i.e., have seemingly random sequence similarity (27). Thus, if we could safely lower the threshold just a little, we would gain many annotations of structural and functional similarity. In fact, any push a little lower reveals many proteins with similar phenotype, e.g., structure or function. Unfortunately, without improving the search method, such a lowering usually comes at the expense of even more proteins with dissimilar phenotype.

This simple reality has been driving the advance of methods using sequence similarity to establish relations: from advanced pairwise comparisons (14,31) over sequence-profile (32-35) to profile-profile comparisons (24,36-41) or efficient shortcuts to the latter (10,42). All those methods share one simple idea, namely, to use evolutionary information (EI) to create families of related proteins. These are summarized in <u>multiple sequence alignments (MSAs)</u>. Using such information as input to machine learning methods has been generating essentially all <u>state-of-the-art (SOTA)</u> prediction methods for almost three decades (43-45). Using MSAs has also been one major key behind the breakthrough in protein structure prediction through *AlphaFold2* (46), and subsequently of *RoseTTAFold* (47) which builds on ideas introduced by *AlphaFold2*, i.e., allowing for communication between different sequence- and structure modules within the network. Transfer- or representation-learning offer a novel route toward comparisons of and predictions for single sequences without MSAs.

### Embeddings capture language of life written in proteins

The introduction of LSTM- or attention-based <u>Language Models (LMs)</u> such as ELMo (48) or BERT (17) enabled a better use of large, unlabeled text corpora which arguably improved all tasks in <u>natural language processing (NLP)</u> (49). These advances have been transferred to proteins through <u>protein Language Models (pLMs)</u> equating amino acids with words in NLP and the sequence of entire proteins with sentences. Such pLMs learn to predict masked or missing amino acids using large databases of raw protein sequences as input (9,12,50-54), or by refining the pLM through another supervised task (16,55). Processing the information learned by the pLM, e.g., by using the output of the last hidden layers of the networks forming the pLMs, yields a representation of protein sequences referred to as <u>embeddings</u> (Fig. 1 in (9)). Embeddings have been used successfully as exclusive input to predicting secondary structure and subcellular localization at performance levels almost reaching (12,50,51) or even exceeding (9,56,57) the SOTA using evolutionary information from MSAs as input. Embeddings can even substitute sequence similarity for homology-based annotation transfer (58,59). The power of such embeddings has been increasing with the advance of algorithms and the growth of data (9). The recent advances have shown that a limit to such improvements has not nearly been reached when writing this (22.02.2022).

**Fig. 1:**
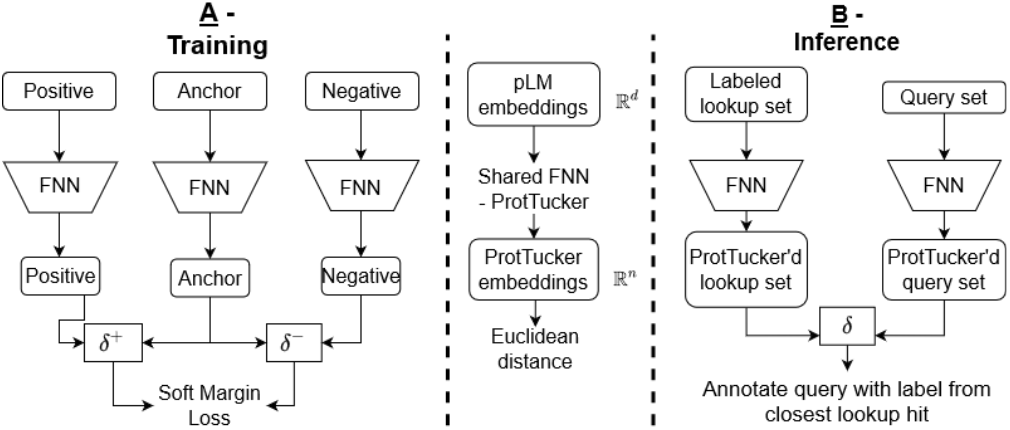
Sketch of ProtTucker. <u>Panel A</u> illustrates how protein triplets were used to contrastively learn the CATH hierarchy (1,2). First, protein Language Models (pLMs) were used as static feature encoders to derive embeddings for protein sequences (anchor, positive, negative). The embedding of each protein was processed separately by the same, shared FNN with hard parameter sharing, called <u>ProtTucker</u>. During optimization, the Soft Margin Loss was used to maximize the distance between proteins from different CATH classes (anchor-negative pairs) while minimizing the distance between proteins in the same CATH class (anchor-positive pairs). All four CATH-levels were simultaneously learned by the same FNN. This resulted in a newly, learned CATH-optimized embedding for each protein. <u>Panel B</u> sketches how the contrastive learning FNN is used for prediction of new proteins (inference). For all proteins in a lookup set with experimental annotations (labeled proteins; here the CATH lookup set), as well as for a query protein without experimental annotations (unlabeled proteins) all embeddings are extracted in two steps: (1) extract per-residue embeddings from original pLM and create per-protein embeddings by averaging over protein length. (2) Input those embeddings into the pretrained FNNs, i.e., ProtTucker. Similar to homology-based inference (HBI), predictions are generated by transferring the annotation of the closest hit from the lookup set to the query protein. The embedding-based annotation transfer (EAT) transferred annotations to the hit with the smallest Euclidean distance in *ProtTucker* embedding space.

Embeddings from pLMs capture a diversity of higher-level features of proteins, including various aspects of protein function and structure (9,12,51,58-61). In fact, pLMs such as ProtT5 (9) or ESM-1b (12) capture aspects about protein structure so impressively that inter-residue distances – and consequently 3D structure – can be predicted without using MSAs, even with relatively small (few free parameters) Deep Learning (DL) architectures (62).

Supervised learning directly maps the input to the class output. Instead, contrastive learning (63), optimizes a new embedding space in which similar samples are pushed closer, dissimilar samples farther apart. Contrastive learning relies only on the similarity between pairs (or triplets) of samples instead of on class label. The definition of similarity in embedding rather than sequence space, combined with contrastive learning, offered an alternative to sequence-based protein comparisons. This led us to hypothesize that we might find structurally and functionally consistent sub-groups within protein families from raw sequences. As a proof-of-principle, a rudimentary precursor of this work helped to cluster FunFams (2,59). The benefit of optimizing embeddings specifically for SCOPe fold recognition (64) has recently been shown (16,60,65). Other approaches toward fold recognition deep learn fold-specific motifs (66), pairwise similarity scores (67) or sequence alignments (68). However, most of the top-performing solutions rely on information extracted from MSAs (69) and do not utilize the transfer-learning capabilities offered by recent pLMs.

Here, we expand on the hypothesis that replacing supervised learning by contrastive learning intrinsically fits the hierarchy of CATH (1,2). We propose an approach that marries both, self-supervised pretraining and contrastive learning, by representing protein sequences as embeddings, and using increasing overlap in the CATH hierarchy as a notion of increasing structural similarity to contrastively learn a new embedding space. We used the pLM ProtT5 (9) as static feature encoder (no fine-tuning of the pLM) to retrieve initial embeddings that were then mapped by a feed-forward neural network (FNN) to a new, learned embedding space optimized on CATH through contrastive learning. More specifically, the Soft Margin Loss was used with triplets of proteins (anchor, positive, and negative) to optimize the new embedding space toward maximizing the distance between proteins from different CATH classes (anchor-negative pairs) while minimizing the distance between proteins in the same CATH class (anchor-positive pairs). Triplets of varying structural similarity were used simultaneously to optimize a single, shared network: all four CATH-levels were simultaneously learned by one FNN. The resulting model was called *ProtTucker* and its embeddings were established to identify more distant relations than is possible from sequence alone. One important objective of *ProtTucker* is to study entire functional modules through identifying more distant relations, as found to be crucial for capturing mimicry and hijacking of SARS-CoV-2 (70).

## Methods

### CATH hierarchy

The CATH (2,22) hierarchy (v4.3) classifies three-dimensional (3D) protein structures from the PDB (Protein Data Bank (71)) at the four levels Class, Architecture, Topology and Homologous superfamily. On average, higher levels (further away from root: H>T>A >C) are more similar in their 3D structure or have more residues for which the same level of 3D similarity is reached. We used increasing overlap in this hierarchical classification as a proxy to define increasing structural similarity between protein pairs. For example, we assumed that any two proteins with the same topology (T) are structurally more similar than any two proteins with identical architecture (A) but different topology (T). In more formal terms: SIM3D(P1,P2)>SIM3D(P3,P4), where T(P1)=T(P2) & T(P3)≠T(P4) & A(P3)=A(P4). This notion of similarity was applied on all four levels of CATH.

### Data set

The sequence-unique datasets provided by CATH (1,2) v4.3 (123k proteins, CATH-S100) provided training and evaluation data for ProtTucker. A test set (300 proteins, dubbed test300 in the following) for final evaluation and a validation set (200 proteins, dubbed ***<u>val200</u>***) for early stopping were randomly split off from CATH-S100 while ensuring that (1) every homologous superfamily appeared maximally once in test300 ⋂ val200 and (2) each protein in test300 & val200 has a so called Structural Sub-group (SSG) annotation, i.e., clusters of domain structure relatives that superpose within 5Å (0.5nm), in CATH. To create the training set, we removed any protein from CATH-S100 that shared more than 20% pairwise sequence identity (PIDE) to any validation or test protein according to MMSeqs2 (10) applying its iterative profile-search (*--num-iterations 3*) with highest sensitivity (*-s 7.5*) and bidirectional coverage (*--cov-mode 0*). Additionally, large families (>100 members) within CATH-S100 were clustered at 95% PIDE and length coverage of 95% of both proteins using MMSeqs2 (bidirectional coverage; *--cov-mode 0*). The cluster representatives were used for training (66k proteins, dubbed ***<u>train66k</u>***) and as lookup set during early stopping on set *val200*. We needed a lookup set from which to transfer annotations because contrastive learning outputs embeddings instead of class predictions. For the final evaluation on *test300*, we created another lookup set but ignored val200 proteins during redundancy reduction (69k proteins, dubbed ***<u>lookup69k</u>***). This provided a set of proteins for validation which had similar sequence properties to those during the final evaluation while “hiding” them during training and not using them for any other optimization. To ensure strict non-redundancy between lookup69k and test300, we further removed any protein from *test300* with HVAL>0 (26) to any protein in *lookup69k* (219 proteins, dubbed ***<u>test219</u>*** in the following). All performance measures were computed using *test219*.

Data augmentation can be crucial for contrastive learning to reach performance in other fields (72). However, no straightforward way exists to augment protein sequences as randomly changing sequences very likely alters or even destroys protein structure and function. Therefore, we decided to use homology-based inference (HBI) for data augmentation during training, i.e., we created a new training set based on Gene3D (5) (v21.0.1) which uses Hidden Markov Models (HMMs) derived from CATH domain structures to transfer annotations from labeled CATH to unlabeled UniProt. We first clustered the 61M protein sequences in Gene3D at 50% PIDE and 80% coverage of both proteins (bidirectional coverage; *--cov-mode 0*) and then applied the same MMSeqs2 profile-search (*--num-iterations 3 –s 7.5*) as outlined above to remove cluster representatives with ≥20% PIDE to any protein in test300 or val200 (PIDE(P_train_,P_test300|val200_)≤20%). This filtering yielded 11M sequences for an alternative training set (dubbed ***<u>train11M</u>***).

The CATH detection of ProtTucker was further analyzed using a strictly non-redundant, high-quality dataset. This set was created by first clustering CATH v4.3 at 30% using HMM profiles from HMMER and additionally discarding all proteins without equivalent entry in SCOPe, i.e., the domain boundaries and the domain-superfamily assignment had to be nearly identical (3186 proteins, CATH-S30). We used the highly sensitive structural alignment scoring tool SSAP (7,8) to compute the structural similarity between all protein pairs in this set.

We probed whether ProtTucker embeddings might also help in solving tasks unrelated to protein structure/CATH, using as proxy a dataset assessing subcellular location prediction in ten states (56,73). We embedding-transferred annotations (EAT) from the standard *DeepLocTrain* set to 490 proteins in a recently proposed test set (*setHard*) that was strictly non-redundant to *DeepLocTrain*. Datasets described elsewhere in more detail (56,73). Finally, we showcased predictions for entire organisms using three UniProt reference proteome: *Escherichia coli* (E. Coli; reviewed, Swiss-Prot (74)), *Armillaria ostoyae* (A. ostoyae; unreviewed, TrEMBL (74)) and *Megavirus Chilensis* (M. Chilensis; unreviewed TrEMBL (74)).

### Data representation

Protein sequences were encoded through distributed vector representations (embeddings) derived from four different pre-trained protein language models (<u>pLM</u>): (1) **<u>ProtBERT</u>** (9) based on the NLP (Natural Language Processing) algorithm BERT (17) but trained on BFD (Big Fantastic Database) with over 2.3 billion protein sequences (11). (2) **<u>ESM-1b</u>** (12) is similar to (Prot)BERT but trained on UniRef50 (74). (3) **<u>ProtT5-XL-U50</u>** (9) (*ProtT5* for simplicity) based on the NLP sequence-to-sequence model T5 (18) trained on BFD and fine-tuned on Uniref50. (4) **<u>ProSE</u>** (16) trained long short-term memory cells (LSTMs) either solely on 76M unlabeled sequences from UniRef90 (**<u>ProSE-DLM</u>**) or on additionally predicting intra-residue contacts and structural similarity from 28k SCOPe proteins (64)(multi-task: **<u>ProSE-MT</u>**). While ProSE, ProtBert and ESM-1b were trained on reconstructing corrupted tokens/amino acids from non-corrupted (protein) sequence context (masked language modeling), ProtT5 was trained by *teacher forcing*, i.e., input and targets were fed to the model with inputs being corrupted protein sequences and targets being identical to inputs but shifted to the right (span generation with span size of 1 for ProtT5). Except for ProSE-MT, all pLMs were optimized only through self-supervised learning exclusively using unlabeled sequences for pre-training.

pLMs output a single vector for each residue yielding an *LxN*-dimensional matrix (L: protein length, N: embedding dimension; N=1024 for ProtBERT/ProtT5; N=1280 for ESM-1b; N=6165 for ProSE). From this *L* x *N* embedding matrix, we derived a fixed-size N-dimensional vector representing each protein by averaging over protein length (Fig. 1 (9)). The pLMs were used as static feature encoder only, i.e., no gradient was backpropagated for fine-tuning. As recommended in the original publication (9), for ProtT5, we only used the encoder part of ProtT5 in half-precision to embed protein sequences. Similarly, ProtBERT embeddings were derived in half-precision.

### Contrastive learning: architecture

A two-layer feedforward neural network (FNN) projected fixed-size per-protein (sentence-level) embed-dings from 1024-d (1280-d/6165-d for ESM-1b and ProSE respectively) to 256 and further to 128 dimensions with the standard hyperbolic tangent (*tanh*) as non-linearity between layers. We also experimented with deeper/more sophisticated networks without any gain from more free parameters (data not shown). This confirmed previous findings that simple networks suffice when inputting advanced embeddings (9,12,62,75). As the network was trained using contrastive learning, no final classification layer was needed. Instead, the 128-dimensional output space was optimized directly.

### Contrastive learning: training

During training, the new embedding space spanned by the output of the FNN was optimized to capture structural similarity using triplets of protein embeddings. Each triplet had an anchor, a positive and a negative. In each epoch, all *train66k* proteins were used once as anchor, while positives and negatives were sampled randomly from *train66k* using the following *<u>hierarchy-sampling</u>*. First, a random level α (α=[1,2,3,4]) describing the increasing structural overlap between triplets was picked. This defined a positive (same CATH-number as anchor up to level α’**≤**α) and a negative label (same CATH-number as anchor up to level α’**<**α). For instance, assume the anchor has the CATH-label 1.25.10.60 (Rad61, Wapl domain) and we randomly picked α=3 (topology-level), only proteins with the anchor’s topology (1.25.10.x; Leucine-rich Repeat Variant) qualify as positives while all negatives share the anchor’s architecture (1.25.y.z; Alpha Horseshoe) with different topology (y≠10). Self-hits of the anchor were excluded. From the training proteins fulfilling those constraints, one positive and one negative were picked at random. If no triplets could be formed (e.g., α=4 with a single-member homologous superfamily had no positive for this anchor/α combination), α was changed at random until a valid triplet could be formed (eventually, all proteins found a class-level partner). This flexibility in sampling enabled training on all proteins, independent of family size.

Unlike randomly sampling negatives, the hierarchical sampling could be described as semi-hard sampling as it zoomed into triplets that were neither too easy (little signal) nor too hard (outliers) to classify by ensuring a minimal overlap between the anchor and the chosen negative/positive pair. Thereby, trivial triplets are under-sampled (avoided), i.e., those with 3D structures so different that the separation becomes trivial (*daylight zone*). As the final triplet selection was still random, anchor-positive pairs could still be too easy/similar which was shown to hinder the success of contrastive learning (76). To solve this issue, we paired hierarchy-sampling with so called *<u>batch-hard sampling</u>* (76) which offers a computationally efficient solution for sampling *semi-hard* triplets within one mini-batch. More specifically, we combined *batch-hard sampling* with the triplets created using *hierarchy-sampling* by re-wiring all proteins, irrespective of anchor, positive or negative, within one mini-batch such that they satisfied the hierarchy-sampling criterion and had maximum/minimal Euclidean distance for anchor-positive/anchor-negative pairs. Sampling hard triplets only within each mini-batch instead of across the entire data set avoided extreme outliers (potentially too hard/noisy) while increasing the rate of semi-hard anchor-positive/anchor-negative pairs. Assume multiple proteins of the topology Leucine-rich Repeat Variant were within one mini-batch, the hardest positive for each anchor would be picked by choosing the anchor-positive pair with the largest Euclidean distance. Accordingly, anchor-negative pairs would be picked based on the smallest Euclidean distance. For each mini-batch, this sampling was applied to all four levels of the CATH-hierarchy, so triplets were re-wired on all four CATH levels resulting in a total batch-size of about: batch_size * 3 * 4. This was an “about” instead of “equal” because for some mini-batches, not all proteins had valid triplets for all four levels.

Finally, the same two-layer FNN was used (hard parameter sharing) to project the 1024-d (or 1280-d/6165-d for ESM-1b or ProSE respectively) embeddings of all proteins, irrespective of anchor, positive or negative, to a new 128-d vector space. The *Soft Margin Loss* was used to optimize this new embedding space such that anchor-positive pairs were pulled together (reduction of Euclidean distance) while pushing apart anchor-negative pairs (increase of Euclidean distance). The efficiency of combining the Soft Margin Loss with batch-hard sampling was established before (76), although without prior hierarchical triplet sampling. Here, we used Soft Margin Loss as implemented in PyTorch:

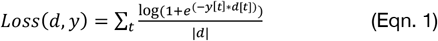

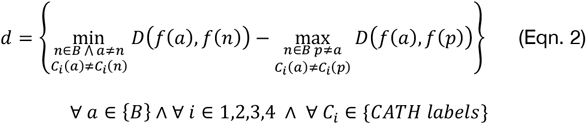

*B* represents one mini-batch created through hierarchical sampling, *f*(*a*), *f*(*p*) and *f*(*n*) represent the *ProtTucker* embeddings of proteins *a* (anchor), *n* (negative), and *p* (positive) represented as pLM embeddings; *C_i_* represents the CATH annotation of a protein on the *i*’th hierarchy level of CATH; finally, D(f(a),f(x)) represents the Euclidean distance between the embeddings for proteins a and x. We created the mini-batch *B* used for training by choosing for each protein or anchor *a* in *B* the hardest negative *n* and the hardest positive *p* by picking those proteins in *B* that have the smallest | largest Euclidean distance *D* to *a ProtTucker* embedding space while not sharing | sharing *C_i_*, respectively. Those semi-hard triplets are indexed by *t* and *d[t]* referring to the difference between *D* of anchor-negative and *D* of anchor-positive. In our case, the label for the *t*’th triplet *y[t]* is always 1 as the sign of x indicates training success, i.e., whether the distance anchor-positive is smaller than that between anchor-negative.

Consequently, triplets of varying structural similarity were used simultaneously to optimize a single, shared network, i.e., all four CATH-levels were learned by the same network at the same time (Fig. 1A). We used the *Adam optimizer* (77) with a learning rate of 0.001, and a batch-size of 256 to optimize the network. The effective batch-size increased due to batch-hard sampling to a maximum of 3072, depending on the number of valid triplets that could be formed within the current mini-batch. Training terminated (*early stopping*) at the highest accuracy in predicting the correct homologous superfamily for set *val200*.

### Evaluation and prediction (inference)

While supervised training directly outputs class predictions, contrastive learning, outputs a new embedding space. Thus, predictions (inferences) were generated as for homology-based inference (HBI), i.e., given protein X with experimental annotation (CATH assignment) and a query protein Q without, then HBI transfers the annotation from X to Q if sequence similarity SIM(X,Q)> threshold. For contrastive learning, we replaced SIM(X,Q) by D(f(X),f(Q)) with D as the shortest Euclidean distance in embedding space (Fig. 1B). In previous studies (9,58,59), we found the Euclidean distance performed better than the cosine distance which is more common in AI/NLP. The Euclidean distance also optimized the ProtTucker embeddings. Set *test219* with the *lookup69k* as lookup set (set of all X) served as final evaluation. If no protein in the lookup set shared the annotation of the query protein at a certain CATH-level (more likely for H than for C), the sample was excluded from the evaluation of this CATH-level as no correct prediction was possible (Table S1).

During evaluation, we compared the performance of our embedding-based annotation transfer (EAT) to HBI using the sequence comparisons from MMSeqs2 (10). While transferring only the HBI hit with the lowest E-value, we searched for hits up to an E-value of 10 to ensure that most proteins had at least one hit while using the highest sensitivity setting (-s 7.5). Additionally, we used publicly available CATH-Gene3D (2) Hidden Markov Models (HMMs) along with HMMER (6) to detect remote homologs up to an E-value of 10.

For both approaches, EAT and HBI, we computed the accuracy as the fraction of correct hits for each CATH-level. A hit at lower CATH-levels could be correct if and only if all previous levels were correctly predicted. Due to varying number of samples at different CATH-levels (Table S1), performance measures not normalized by background numbers could be higher for lower levels. Predictions were counted as incorrect if a query did not have a hit in the lookup set but a lookup protein of the same CATH annotation existed. This not only affected the number of proteins available at different CATH-levels (Table S2) but also the number of classes (Table S3). A random baseline was computed by transferring annotations from a randomly picked protein in *lookup69k* to *test219*.

### Performance measures

The four coarse-grained classes at the top CATH level (“C”) are defined by their secondary structure content. These four branch into 5481 different superfamilies with distinct structural and functional aspects (CATH v4.3.0). However, most standard metrics are defined for binary cases which requires some grouping of predictions into four cases: 1) TP (true positives): correctly predicted to be in the *positive* class, 2) TN (true negatives): correctly predicted to be in the *negative* class, 3) FP (false positives): incorrectly predicted to be positives, and 4) FN (false negatives): incorrectly predicted to be in in the negative class. Here, we focused on performance measures applicable for multiclass problems and are implemented in scikit (78). These were in particular: **<u>accuracy</u>** (Acc, Eqn. 3) as the fraction of correct predictions

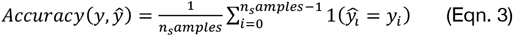

with *y_i_* being the ground truth (experimental annotation) and 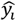 the prediction for protein *i*. In analogy, we defined **<u>coverage</u>** as the proportion of the *test219* proteins for which a classifier made a prediction at a given prediction reliability 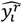 and reliability threshold *θ*:

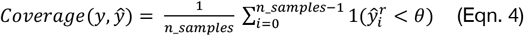

In these definitions accuracy corresponds to precision, and coverage to recall binarizing a multiclass problem through micro-averaging, i.e., by counting the total TPs, FPs and FNs globally, irrespective of the class. The multi-class extension of **Matthew’s correlation coefficient (MCC**, (45)) was defined as:

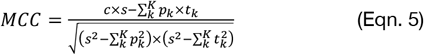

with 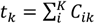 as the number of times class *k* truly occurred, 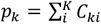 as the number of times class *k* was predicted, 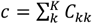, the total number of samples correctly predicted, and 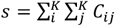, the total number of samples.

**<u>95% confidence intervals</u>** for accuracy and MCC were estimated over n=1000 bootstrap sets; for each bootstrap set we randomly sampled predictions from the original test set with replacement. Standard deviation (or in the case of bootstrapping: bootstrap standard error) was calculated as the difference of each test set (*x_i_*) from the average performance 〈*X*〉 (Eqn. 6). 95% confidence intervals were estimated by multiplying the bootstrap standard error by 1.96.

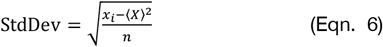

## Results

### Generalization of HBI to EAT

<u>Homology-based inference (HBI)</u> uses sequence similarity to transfer annotations from experimentally characterized (labelled) to uncharacterized (unlabeled) proteins. More specifically, an unlabeled query protein Q is aligned against a set of proteins X with experimental annotations (dubbed *lookup set*) and the annotation of the best hit, e.g., measured as lowest E-value, is transferred if it is below a certain threshold (e.g., E-value(Q,X)<10-3). This relates to inferring the annotation of a query protein from the k-Nearest Neighbors (k-NN) in sequence space. More recently, similar approaches expanded from distance in sequence to distance in embedding space (Fig. 1B) as means for k-NN based annotation transfer (58,60). Here, we refer to such methods as <u>embedding-</u> <u>based annotation transfer (EAT)</u>. We used EAT as a proxy for the comparison of embeddings from five different <u>protein Language Models (pLMs)</u>: ProSE-DLM & ProSE-MT (16), ProtBERT & ProtT5 (9), and ESM-1b (12). Next, we used triplets of proteins (anchor, positive, negative) to learn a new embedding space by pulling protein pairs from the same CATH class (anchor-positive) closer together while pushing apart pairs from different CATH classes (anchor-negative; Fig. 1A). We referred to this method as *ProtTucker*. At this stage, we did not fine-tune the pre-trained pLMs. Instead, we created a new embedding space using a two-layer feed-forward neural network (FNN).

### EAT with raw embeddings level with HBI

First, we transferred annotations from all proteins in *lookup69k* to any protein in *test219*. All pLMs significantly (at 95% CI - confidence interval) outperformed random annotation transfer (Table 1). Performance differed between pLMs (Table 1), with ProtBERT (9) being consistently worse than LSTM-based ProSE-DLM or more advanced transformers (ESM-1b, ProtT5). ESM-1b and ProtT5 also numerically outperformed ProSE-DLM and HBI using MMseqs2 (10), especially on the most fine-grained and thus hardest level of superfamilies. However, MMseqs2 had been used for redundancy-reduction, i.e., the data set had been optimized for minimal performance of MMseqs2. HBI using publicly available HMM-profiles from CATH-Gene3D (2) along with the profile-based advanced HMMER (6) designed for more remote homology detection, outperformed all raw embeddings for homologous superfamilies, while embeddings from ESM-1b and ProtT5 appeared superior on the class- and architecture-level (Table 1). In fact, all HBI values, for MMseqs2 and HMMER, were highest for the H-level, and 2^nd^ highest for the C-level. In contrast, raw pLM embeddings mirrored the random baseline trend, with numbers being inversely proportional to the rank in C, A, T, H (highest for C, lowest for H, Table 1).

**Table 1:** Accuracy for annotation transfer to queries in test219 *_**_.

|  | Method/Input | C | A | T | H | Mean |
| --- | --- | --- | --- | --- | --- | --- |
| Baseline | Random | 29 ±6 | 9 ±4 | 1 ±2 | 0 ±0 | 10 ±3 |
| HBI | MMSeqs2 (sequence) | 52 ±7 | 36 ±6 | 29 ±6 | 35 ±6 | 38 ±6 |
|  | HMMER (profile) | 70 ±6 | 60 ±6 | 59 ±7 | 77 ±7 | 67 ±6 |
| EAT - Unsupervised | ProSE-DLM | 74 ±6 | 48 ±7 | 28 ±6 | 25 ±7 | 44 ±6 |
|  | ESM-1b | 79 ±5 | 61 ±6 | 50 ±7 | 57 ±8 | 62 ±7 |
|  | ProtBERT | 67 ±6 | 38 ±6 | 22 ±6 | 18 ±6 | 36 ±6 |
|  | ProtT5 | 84 ±5 | 67 ±6 | 57 ±6 | 64 ±8 | 68 ±6 |
| EAT - Supervised | ProSE-MT | 82 ±5 | 65 ±6 | 52 ±7 | 56 ±8 | 64 ±7 |
| EAT - Contrastive learning – ProtTucker | ProSE-DLM | 78 ±4 | 53 ±6 | 32 ±6 | 29 ±7 | 48 ±6 |
|  | ProSE-MT | 87 ±4 | 68 ±6 | 53 ±7 | 55 ±8 | 66 ±6 |
|  | ESM-11b | 87 ±4 | 68 ±6 | 59 ±7 | 70 ±7 | 71 ±6 |
|  | ProtBERT | 81 ±5 | 52 ±7 | 37 ±6 | 39 ±8 | 52 ±7 |
|  | ProtT5 | **89 ±4** | 75 ±6 | 64 ±6 | 76 ±6 | 76 ±6 |
|  | ProtT5 (train11M) | 88 ±4 | **77 ±5** | **68 ±5** | **79 ±7** | **78 ±6** |

\* Accuracy (Eqn. 3) for predicting CATH (2,3) levels (C, A, T, H) by transferring annotations from *Lookup69k* (lookup set) to *test219* (queries); shown for each of the four levels from the most coarse-grained level class C to the most fine-grained level of homology H. The column *Mean* marked the simple arithmetic average over the four performance values. Queries with at least one lookup protein of the same CATH classification but without any hit at E-value<10 for MMSeqs/HMMER were counted as incorrect predictions. Errors indicate bootstrapped 95% confidence intervals, i.e., 1.96 bootstrap standard errors (Eqn. 6). Queries with at least one lookup protein of the same CATH annotation but without any hit (no hit with E-value<10 for MMSeqs/HMMER; irrelevant for EAT) were counted as wrong predictions. **Bold** letters mark the numerically highest values (averages over all *test219* proteins) in each column irrespective of the confidence interval.
**Methods:** Baseline: Random transferred the label of a randomly picked protein; HBI: MMSeqs2 (10) used single sequence search to transfer the annotation of the hit with the lowest E-value; HBI: HMMER used HMM-profiles (6); EAT-unsupervised: embedding-based transfer of annotations using the smallest Euclidean distance measured in embedding space of unsupervised pLMs ProSE-DLM, ESM-1b (12), ProtBERT and ProtT5 (9); EAT-supervised: annotation transfer using ProSE-MT trained on structural data in SCOPe; EAT: contrastive learning ProtTucker: contrastive learning trained on CATH classifications in *train66k* using as input embeddings from ProSE-DLM, ProSE-MT, ESM-1b, ProtBERT, and ProtT5; ProtTucker-ProtT5 (train11M) trained on additional data from Gene3D (*train11M*).

### EAT improved through supervised embeddings

ProSE-MT expands ProSE-DLM by additionally training on intra-residue contacts and structural similarity using labeled data from SCOPe (16). This additional effort was reflected by the higher performance for all CATH levels (Table 1, ProSE-MT>ProSE-DLM). The supervision pushed the LSTM-based ProSE-MT to reach performance levels close to the unsupervised, raw embeddings from transformer-based ProtT5. The performance gap increased with classification difficulty (Table 1, ProtT5>ProSE-MT, especially at the H-level).

### EAT improved by contrastively learning embeddings

Contrastive learning tries to bring members from the same class/CATH-level closer while pushing those from different classes further apart. One success is the degree to which these two distributions (same vs. different) were separated through training: the distribution of all pairwise Euclidean distances within (intra/same) and between (inter/different) superfamilies in *train66k* changed substantially through contrastive learning (Fig. 2). Before applying contrastive learning, the distributions between (inter, Fig. 2: red) and within (intra, Fig. 2: blue) overlapped much more than after.

**Fig. 2:**
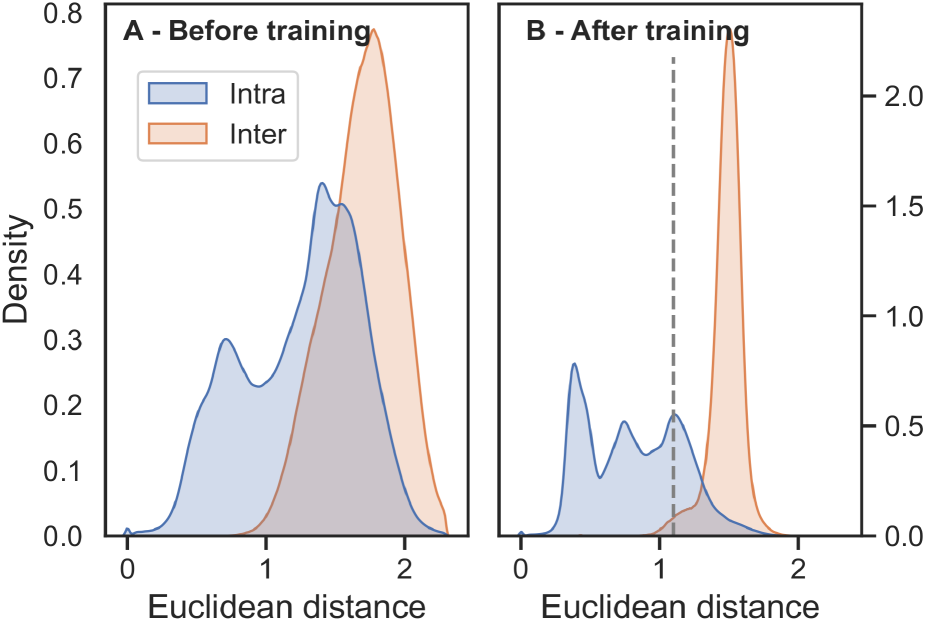
Contrastive learning separated positives from negatives. The structural similarity defined by increasing overlap in CATH drove the contrastive learning of a new embedding space. The new embeddings distanced protein pairs with different CATH classifications (red; same topology but different superfamily) while focusing pairs with the same CATH classification (blue; same superfamily). These graphs compared the Euclidean distance for all such pairs from the set *train66k* before (<u>Panel A</u>) and after (<u>Panel B</u>) contrastive training. Input to the FNN were the raw embeddings from ProtT5 (9), output were the new ProtTucker(ProtT5) embeddings. The dashed line at Euclidean distance of 1.1 in B marked the threshold at which EAT performances started to decrease (Fig. 5).

Displaying the information learned by the embeddings, we compared t-SNE projections colored by the four main CATH classes before (Fig. 3A) and after (Fig. 3C) contrastive learning. These two projections compared 1024 dimensions from ProtT5 (Fig. 3A) with 128 dimensions from ProtTucker (Fig. 3C). To rule out visual effects from higher dimensionality, we also compared the untrained, randomly initialized version of ProtTucker using pre-trained ProtT5 embeddings as input (Fig. 3B). For all cases, the data set (*train66k*) and the parameters for dimensionality reduction were identical. T-SNE projections of raw ProtT5 embeddings qualitatively suggested some class separation (clustering). The information underlying this separation was preserved when projecting ProtT5 embeddings through an untrained ProtTucker (Fig. 3B). Embeddings from ProtTucker(ProtT5), i.e., those resulting through contrastive learning, separated the classes much more clearly.

**Fig. 3:**
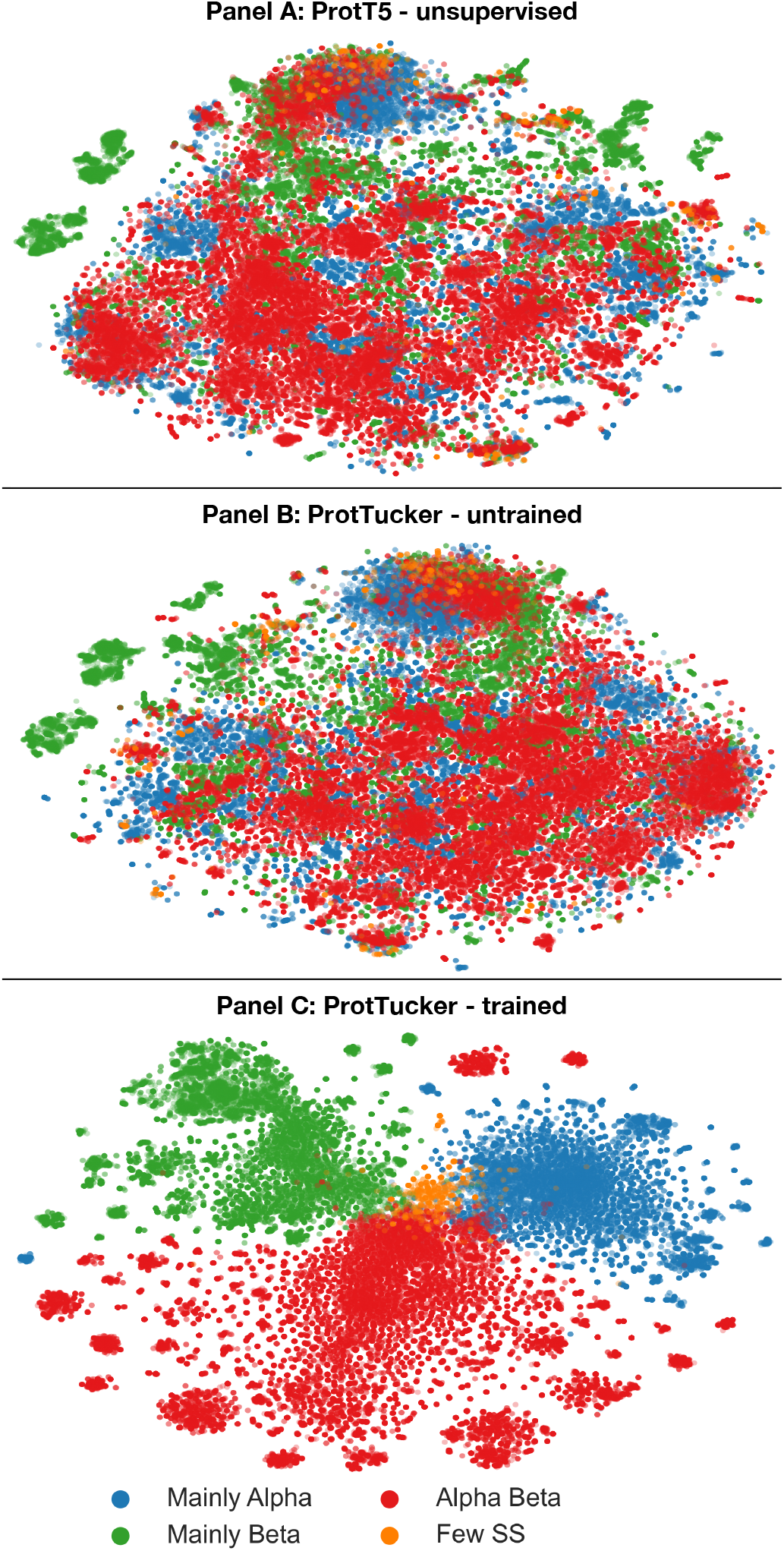
Better CATH class-level clustering. Using t-SNE (4), we projected the high-dimensional ProtTucker(ProtT5) embedding space onto 2D *before* (<u>Panel A</u>; ProtT5) and *after* (<u>Panel C</u>; ProtTucker(ProtT5)) contrastive learning. <u>Panel B</u> visualized the same data embedded with an untrained version of ProtTucker to assess the impact of different embedding dimensions (1024-d for ProtT5 vs. 128-d for ProtTucker(ProtT5)). For all plots, dimensionality was first reduced by Principal Component Analysis (PCA) to 50 dimensions and parameters of the subsequent t-SNE were identical (perplexity=150, learning_rate=400, n_iter=1000, seed=42). The colors mark the major class level of CATH (C), distinguishing proteins according to their major distinction in secondary structure content.

To further probe the extent to which contrastive learning captured remote homologs, we compared the Euclidean distance between all protein pairs in a 30% non-redundant dataset (CATH-S30) with the structural similarity of those pairs computed via SSAP (7,8) (Fig. 4). From the ~10M possible pairs between the 3,186 proteins in CATH-S30 (problem not fully symmetric, therefore N*(N-1): 10.1M), 7.1M had to be discarded due to low quality (SSAP-score <50), leaving 2.9M pairs of which only 1.8% (53k pairs) had the same homologous superfamily (Fig. 4: blue). Despite this imbalance, unsupervised ProtT5 (Fig. 4A) already to some extent separated the same from different superfamilies. Still, ProtTucker(ProtT5) improved this separation, especially, for pairs with low structural similarity (Fig. 4B). This was supported by the Spearman correlation coefficient between the structural similarity and the Euclidean distance increasing from 0.05 to 0.22 after contrastive learning. When considering only the subset of pairs that likely have similar folds (SSAP-score>70), this correlation increased to 0.26 and 0.37 for ProtT5 and ProtTucker(ProtT5), respectively.

**Fig. 4:**
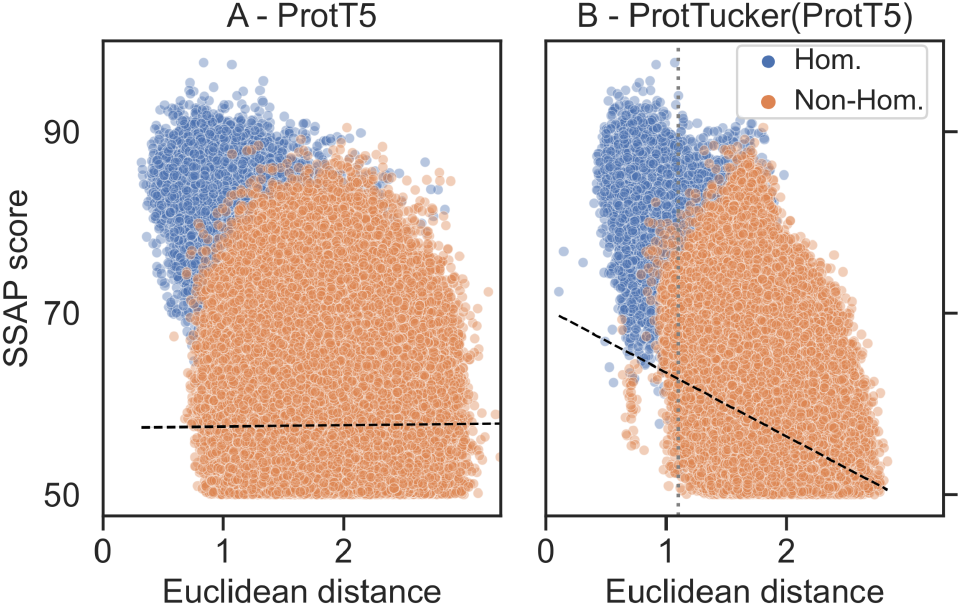
ProtTucker captured fine-grained structural similarity. 3186 non-redundant proteins (CATH-S30) probed the remote homology detection of embeddings *before* (<u>Panel A</u>) and *after* contrastive learning (Panel B). The Euclidean distance between ProtTucker embeddings (<u>Panel B</u>) correlated better with structural similarity computed by SSAP (7,8) than unsupervised embeddings (Panel A): Spearman ρ=0.22 and ρ=0.05 (black dashed lines). This correlation increased to ρ=0.37 and ρ=0.26 for structurally more similar protein pairs (SSAP-score>70). Only 1.8% (53k) of all structurally similar pairs were in the same homologous superfamily (blue). The unsupervised ProtT5 already separated homologous pairs from others, but ProtTucker(ProtT5) improved, especially, for hard cases with low structural similarity. The gray dashed line at Euclidean distance=1.1 in *Panel B* marked the threshold at which EAT performances started to decrease (Fig. 5).

The trend captured by the better separation of distributions (Fig. 2) and structural features (Fig. 3, Fig. 4) translated directly into performance increases: all embeddings optimized on the CATH hierarchy through contrastive learning yielded better EAT classifications than the raw embeddings from pre-trained pLMs (Table 1). As ProtTucker described the process of refining raw embeddings through contrastive learning, we used the annotation <u>ProtTucker(X)</u> - in this section also shortened to <u>PT(X)</u> - to refer to the embeddings output by inputting the pre-trained pLM *X* into the contrastive learning. The improvements were larger for more fine-grained CATH levels: all models improved significantly for the H-level, while only PT(ProtBERT) and PT(ESM-1b) improved from 4 to 14 or from 0 to 21 percentage points for the C-, and the H-level, respectively. PT(ProtT5) consistently outperformed all other pLMs on all four CATH-levels, with an increasing performance gap toward the more fine-grained H-level at which all pLMs except for PT(ESM-1b) performed significantly worse. The improvements from contrastive learning for PT(ProSE-DLM) and PT(ProSE-MT) were mostly consistent but largely insignificant. Especially, the model already optimized using labeled data (ProSE-MT) hardly improved through another round of supervision by contrastive learning and even worsened slightly at the H-level.

We augmented the training set for PT(ProtT5) by adding HBI-hits from HMM-profiles provided by CATH-Gene3D (if sequence dissimilar to test300/val200). This increased the training set from 66k (66*10^3^) to 11m (11*10^6^) proteins (15-fold increase) and raised performance, although the higher values were neither statistically significant nor consistent (Table 1: values in last row not always higher than those in 2^nd^ to last row).

### Ablation study

We studied the effect of *batch-hard* and *hierarchical* sampling on performance by removing each component when training PT(ProtT5) (Table 2). Benchmarking on EAT from *lookup69k* to *test219* established that removing either component lowered performance for all CATH levels. Dropping both sampling methods substantially lowered performance. While dropping *batch-hard* sampling still reached high performance for the coarse-grained C- and A-level, dropping hierarchy-sampling dropped performance for both. Dropping both sampling technique, performed worse for all CATH levels but the decrease for the more fine-grained superfamily level was much larger than for the C-level.

**Table 2:** Ablation study. *.

|  | C | A | T | H | Mean |
| --- | --- | --- | --- | --- | --- |
| Baseline | 89 ± 4 | 75 ± 6 | 64 ± 6 | 76 ± 6 | 76 ± 6 |
| -batch-hard | 88 ± 4 | 73 ± 6 | 62 ± 7 | 69 ± 7 | 73 ± 6 |
| -hierarchy | 83 ± 5 | 69 ± 6 | 62 ± 7 | 71 ± 7 | 71 ± 6 |
| -both | 83 ± 5 | 63 ± 6 | 51 ± 7 | 57 ± 8 | 64 ± 7 |

\* **Accuracy** (Eqn. 3) and 95% CI (Eqn. 6) for predicting CATH-levels (2,3) through EAT from *Lookup69k* (lookup set) to *test219* (queries). We investigate the effect on performance when dropping either *batch-hard* sampling, *hierarchy*-sampling or *both* from the *Baseline* model (ProtTucker(ProtT5)).

### Embedding distance correlated with accuracy

The MCC, (Eqn. 5) of HBI inversely correlated with E-value (Fig. 6, HBI-methods): more significant hits (lower E-values) more often shared the same CATH level than less significant hits (higher E-values). In analogy, we explored the corresponding relation for EAT, namely the correlation between accuracy (Eqn. 3) and embedding distance for ProtTucker(ProtT5). Indeed, accuracy correlated with embedding distance (Fig. 5: solid lines) while recall inversely correlated (Fig. 5: dashed lines) for all four classes. For instance, when transferring only annotations for closest hits with Euclidean distances of 1.1 or less, predictions were made for 57%, 57%, 59% or 75% of the test set (coverage, Eqn. 4) of these 96%, 93%, 91% or 90% were correct for levels C, A, T, H, respectively.

**Fig. 5:**
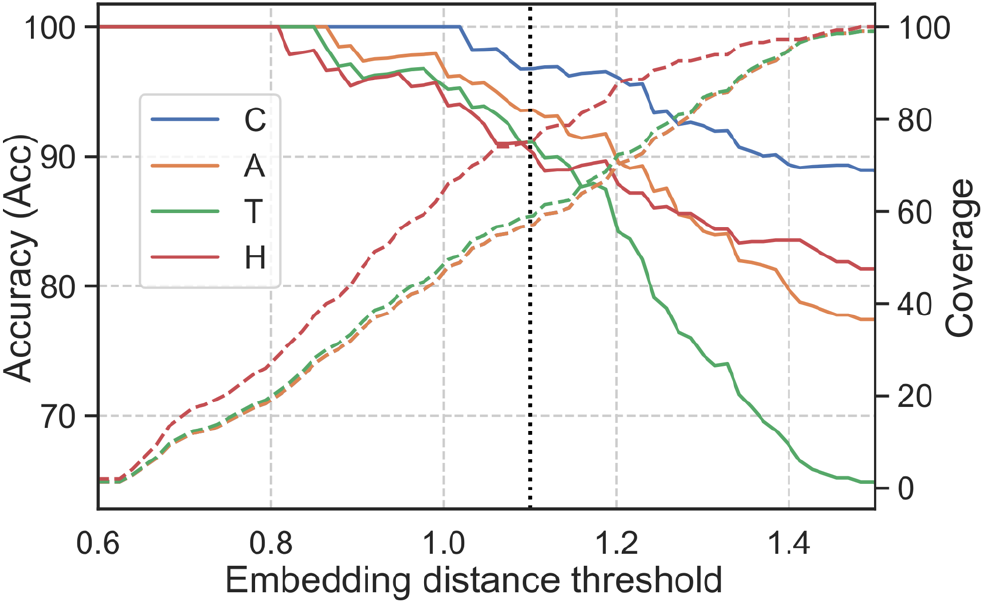
Embedding distance correlated with reliability. Similar to varying E-value cut-offs for HBI, we examined whether the fraction of correct predictions (accuracy; left axis; Eqn. 3) depended on embedding distance (x-axis) for EAT. This was shown by transferring annotations for all four CATH levels (Class: blue; Architecture: orange; Topology: green; Homologous superfamily: red) from *lookup69k* to the queries in set *test219* (Panel B in Fig. 1) using the hit with lowest Euclidean distance. The fraction of *test219* proteins having a hit below a certain distance threshold (coverage, right axis, dashed lines; Eqn. 4) was evaluated separately for each CATH level. For example, at an Euclidean distance of 1.1 (vertical dotted line), 75% of the proteins found a hit at the H-level (Cov(H)=75%) and 90% were correctly predicted (Acc(H)=90%; SOM Tables S3 and S4 for more details). Thus, decreasing embedding distance correlated with EAT performance. This correlation enables users to select only the, e.g., 10% top hits, or as many hits to a certain CATH level as possible, depending on the objectives.

**Fig. 6:**
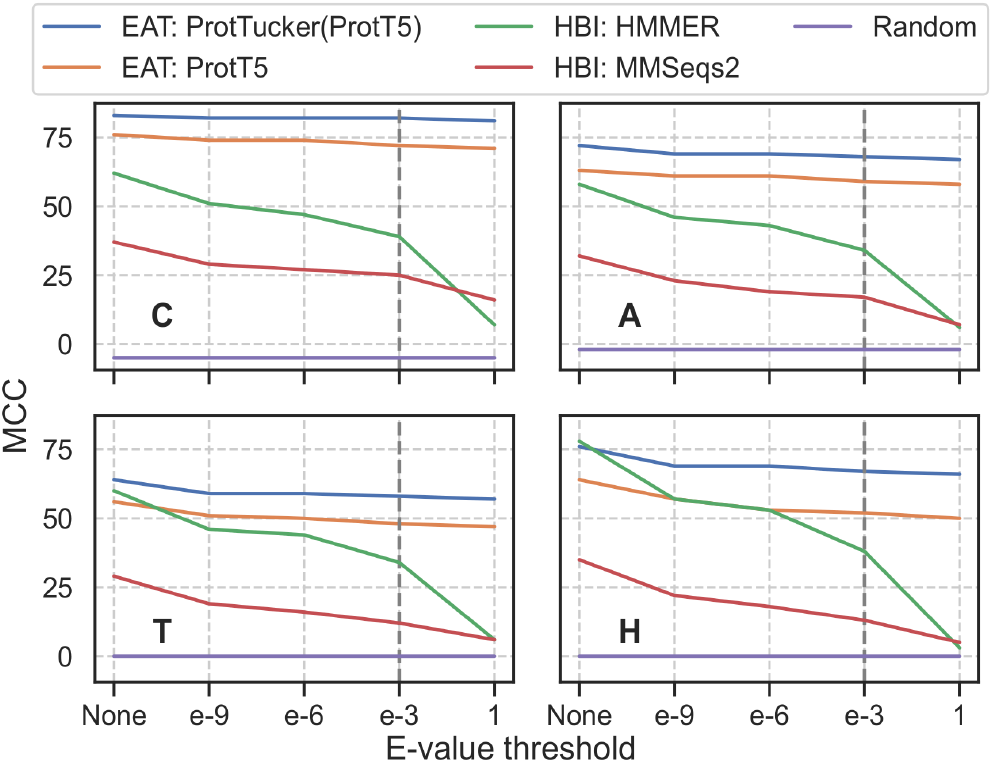
Performance decreasing with lower residual sequence similarity. We analyzed the change of performance in MCC (Eqn. 3) through removing proteins from *lookup69k* based on their E-value with respect to *test219* for two HBI-based (green: HMMER (6); red: MMSeqs2 (10)) and two EAT-based methods (orange: raw ProtT5 (9); blue: contrastive learning optimized ProtTucker(ProtT5)). The Evalues were derived by searching sequences in *test219* against *lookup69k* using (i) HMM-profiles from CATH-Gene3D (2) through HMMer and (ii) MMSeqs2 sequence search with highest sensitivity (*-s 7.5, -cov 0*). “None” referred to the performance without applying any threshold, i.e., all proteins in lookup69k were used for annotation transfer; all other thresholds referred to removing proteins below this E-value from *lookup69k*. Predictions were considered as false positives when no hit was found; pairs without CATH class matches were ignored. While the performance of EAT using raw ProtT5 and refined ProtTucker(ProtT5) embeddings decreased upon removing sequence similar pairs (toward right), HBI-based methods dropped significantly more. The default threshold for most sequence searches (E-value<1e-3) was highlighted by vertical, gray, dashed lines.

### ProtTucker reached into the midnight zone

Annotation transfer by HBI crucially depends on the sequence similarity between query (unknown annotation) and template (experimental annotation). Usually, the significance of an inference is measured as the chance of finding a hit at random for a given database size (E-value; the lower the better). Here, we compared the effect of gradually removing hits depending on their E-values. Essentially, this approach measured how sensitive performance was to the degree of redundancy reduction between query and lookup set. For instance, at a value of 10^-3^ (dashed vertical lines in Fig. 6), all pairs with E-values≤10^-3^ were removed. HBI based on sequence alone performed much better with than without residual redundancy (Fig. 6). The trend was similar for EAT, but much less pronounced: EAT succeeded for pairs with very different sequences (Fig. 6 toward right) almost as well as for proteins with more sequence similar matches in the set (Fig. 6 toward left: EAT almost as high as toward right).

### ProtTucker not a generalist

We evaluated the generality of ProtTucker embeddings by (mis)-using them as exclusive input to predict subcellular location in ten states. To this end, we EAT transferred annotations from an established data set (Table S5) to a strictly non-redundant test set (*setHard*, Table S5). ProtTucker(ProtT5) embeddings outperformed the raw ProtT5 embeddings in the CATH classification for which they were optimized (structural similarity; Table 1), there appeared no performance gain in predicting location. Conversely, performance also did not decrease significantly, indicating that the new embeddings retained some of the information available in ProtT5 embeddings.

### Family size mattered

By clustering very large protein families (>100 members after redundancy reduction) at 95% PIDE, we constrained the redundancy in set *train66k*. Nevertheless, when splitting *test219* into three bins of varying family sizes, we still observed a trend towards higher accuracy (Eqn. 3) for larger families at the H-level (Fig. S1). We chose the three bins such that they contained about the same number of samples (small families: ≤10 members, medium: 11-70, and large: ≥70 members). Especially, unsupervised EAT using the raw ProtT5 embeddings exhibited a clear trend towards higher accuracy with increasing family size. In contrast, the two HBI-methods (MMseqs2, HMMER), as well as EAT using the optimized ProtTucker(ProtT5) embeddings performed similarly for small and medium-sized families and much better for large families.

### EAT complements HBI

As previously shown (59), ProtTucker can improve clustering functional families (79). Here, we showed how EAT can be used to detect outliers. Firstly, we computed pairwise Euclidean distances between the embeddings of all protein pairs in set *train66k* and analyzed the five pairs (10 proteins) with the highest Euclidean distance in the same homologous superfamily (Table S6). High distance within the same homologous superfamily indicates potentially wrong annotations. Secondly, we computed the nearest neighbors of those ten proteins to find an alternative, potentially more suitable annotation. For instance, the proteins in the *Phosphorylase Kinase* superfamily with the largest embedding distance (4pdyA01, bacterial aminoglycoside phospho-transferase) to any other protein within this family (3skjF00, human *Galactose-binding domain-like* (80)) linked to different UniProt entries (*C8WS74_ALIAD* and *EPHA2_HUMAN*). In contrast, the nearest neighbor (3heiA00, human *phosphorylase kinase* (81)) of 3skjF00 linked to the same UniProt entry (*EPHA2_HUMAN* (74)) with the same enzymatic activity (EC number 2.7.10.1 (82)). Such analyses may indicate impure homologous superfamilies along with suggesting alternative labels to be confirmed or rejected through manual curation.

### EAT predicts entire proteomes in minutes

Training ProtTucker(ProtT5) required generating ProtT5 embeddings for *train66k*. This took 23m and 11m, respectively. Embeddings were generated using ProtT5 in half-precision with batch processing. All times were measured on a single Nvidia RTX A6000 with 48GB of vRAM and an AMD EPYC ROME 7352.

When predicting for new queries, ProtTucker requires *labeled lookup* proteins from which annotations can be transferred to unlabeled query proteins. Embeddings for this lookup set are pre-computed for the first query and can be re-used for all subsequent queries at any future time. The time required to labeled *lookup* proteins from which annotations can be transferred to unlabeled query proteins. Embeddings for this lookup set are pre-computed for the first query and can be re-used for all subsequent queries at any future time. The time required to generate ProtTucker embeddings from the embeddings of pLMs was negligible as its generation required only a single forward pass through a two-layer FNN. This implied that the total time for EAT with ProtTucker was largely determined by the embedding generation speed. For instance, creating per-protein embeddings from ProtT5 for the 123k proteins in CATH-S100 required 23 minutes (m). The total time for creating ProtTucker(ProtT5) embeddings from ProtT5 embeddings for the same set on the same machine was 0.5 seconds (s), i.e., ProtTucker added about 0.3%. Creating HMM profiles for the same set using either MSAs from MMSeqs2 (*--num-iterations 3, -s 7.5*) or jackhmmer took 15m or 30h, respectively (Table 3).

**Table 3:** Runtime. *.

| Methods | Pre-processing [s] | Inference [s] |
| --- | --- | --- |
| MMSeqs2 (sequence) | $0.2 \cdot 10^3$ | $2.5 \cdot 10^{-2}$ |
| HMMER (HMM) | $114 \cdot 10^3$ | $150 \cdot 10^{-2}$ |
| ProtTucker(ProtT5) | $1.4 \cdot 10^3$ | $1.4 \cdot 10^{-2}$ |

\* Runtime to transfer annotations from CATH-S100 (123k proteins) to a single query. All times measured in seconds [s] on a single Nvidia RTX A6000 with 48GB of vRAM and an AMD EPYC ROME. **Methods:** two HBI-based methods (MMSeqs2 and HMMER) and one EAT-based method (ProtTucker(ProtT5)). **Pre-processing** measured the time required for building datasets (indexed database for MMSeqs2; MSA for jackhmmmer plus HMM profiles (HMMER) or ProtT5 embeddings (ProtTucker); **Inference** the time for each new protein with a transfer.

To predict using EAT, users have to embed only single query proteins requiring, on average, 0.01s per protein for the CATH-S100 set. Using either single protein sequence search (MMSeqs2), pre-computed HMM profiles (HMMER) or pre computed embeddings (ProtTucker) to transfer annotations from CATH-S100 to a single query protein took on average 0.025s, 1.5s or 0.0008s, respectively. Proteins in the PDB and CATH are, on average, roughly half as long (173 residues) as those from UniProt (343 residues). This is relevant for runtime, because embedding generation scales quadratically with sequence length (Fig. 13 in SOM of (9)).

This increase was also reflected for the proteome-wide annotation transfer (Table 4), although these values included computations required for all aspects of EAT (1: load ProtT5 embeddings for pre-computed CATH-S100 lookup set; 2: load ProtT5 and embedding for query proteome; 3: generate ProtTucker(ProtT5) embeddings for queries and lookup; 4: compute pairwise Euclidean distances between query/lookup). We compared EAT using ProtTucker(ProtT5) embeddings to HBI proxied by existing Gene3D annotations taken from UniProt for three different proteomes (Table 4). At an expected error rate of 5% (Euclidean distance ≤0.9, Table S3), EAT predicted substantially more proteins than Gene3D at HMMER E-value<10-3. For the subset of proteins for which both methods transferred annotations, those largely agreed (*Agreement*, Table 4; Table S7 for other thresholds). All values for coverage decreased for multi-domain proteins, as proxied by “multiple Gene3D annotations”, while the agreement between Gene3D and ProtTucker(ProtT5) increased for 2 of 3 proteomes (Table 4: *multi*).

**Table 4:**
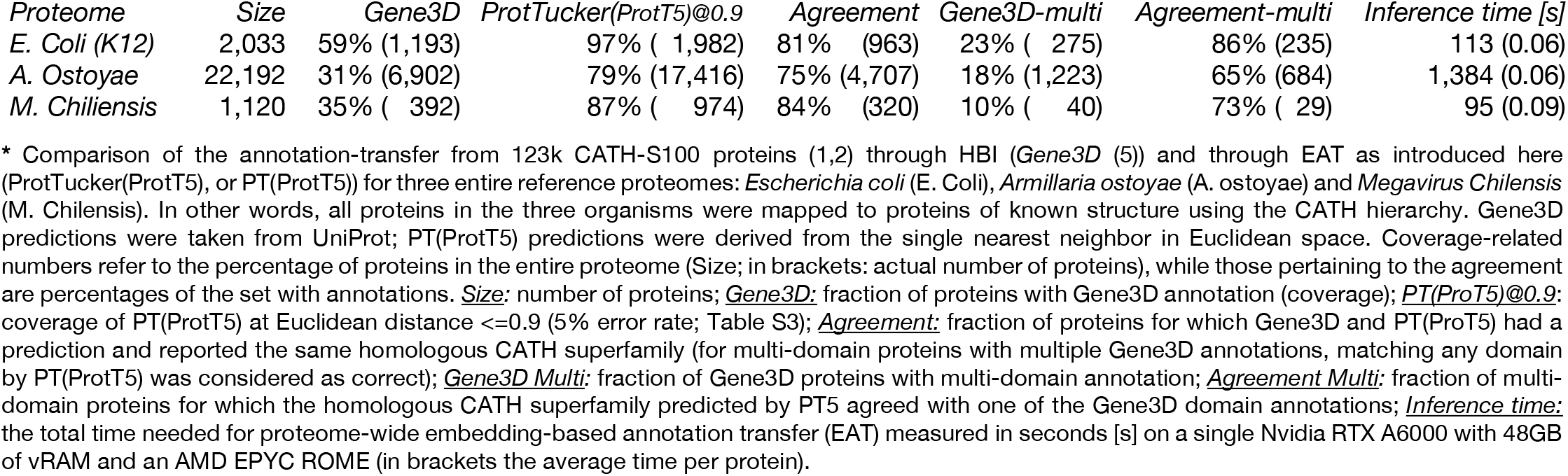
CATH predictions for three model proteomes. *.

## Discussion

### Prototype for representation learning of hierarchies

We have presented a new solution for combining the information implicitly contained in the embeddings from protein Language Models (pLMs) and contrastive learning to learn directly from hierarchically sorted data. As proof-of-concept, we applied the concept to the CATH hierarchy of protein structures (2,3,22,83). Hierarchies are difficult to handle by traditional supervised learning solutions. One *shortcut* is to learn each level in the hierarchy independently (84-86) at the price of having less information for other levels and of not explicitly benefiting from the hierarchy. Instead, our solution of contrastively learning protein triplets (anchor, positive, negative) to extract a new embedding space by condensing positives and moving negatives apart benefits from CATH’s hierarchical structure. Simultaneously training a single, shared feed-forward neural network (FNN) on triplets from all four CATH classification levels allowed the network to directly capture the hierarchy.

Encoding protein sequences through previously trained pLMs enabled ready information transfer from large but unlabeled sequence databases such as BFD (11) to 10,000-times smaller but experimentally annotated (labeled) proteins of known 3D structure classified by CATH. In turn, this allowed us to readily leverage aspects of protein structure captured by pLMs that are informative enough to predict structure from embeddings alone (62). Although the raw, pre-trained, unoptimized embeddings captured some aspects of the classification (Fig. 2A, Fig. 3A, Fig. 4A, Table 1), contrastive learning boosted this signal significantly (Fig. 2B, Fig. 3C, Fig. 4B, Table 1).

Crucial for this success was the novel combination of hierarchy- and batch-hard sampling (Table 2). Presumably, because those techniques enforce so-called *semi-hard* triplets that are neither too simple nor too hard to learn (76). This training setup learned different classifications for the same protein pair, depending on the third protein forming the triplet, thereby forcing the network to learn the complex hierarchy. The ambivalence in the notion of *positive/negative pair* facilitated training by allowing to include superfamilies with few members (otherwise to be skipped) and it increased the number of possible triplets manifold compared to only sampling on the level of superfamilies. These advantages might partially explain the synergy of both sampling techniques (Table 2).

### Raw embedding EAT matched profile alignments in hit detection

In technical analogy to homology-based inference (HBI), we used embedding based annotation transfer (EAT, Fig. 1B) to transfer annotations from labeled lookup proteins (proteins with a known CATH classification) to unlabeled query proteins (any protein of known sequence without known structure). Instead of transferring annotations from the closest hit in sequence space, EAT transferred annotations to the hit with smallest Euclidean distance in embedding space. This relatively simple approach was shown previously to predict protein function as defined by Gene Ontology (GO) better than hand-crafted features (60) even to levels competitive to much more complex approaches (58).

The concept of EAT was so successful that raw embeddings from two different pre-trained pLMs (ESM-1b (12), and ProtT5 (9)) already set the bar high for predicting CATH levels. The raw, general-purpose ESM-1b and ProtT5 outperformed HBI based on advanced HMM-profiles from HMMER (6) on the C- and A-level while falling short on the H-level (Table 1). Furthermore, we showed that ProtT5 already separated protein pairs with the same from those with different homologous superfamilies even when using a lookup set that consisted only of proteins with maximally 30% pairwise sequence identity (Fig 4A). Importantly, this competitive performance was achieved at a much smaller cost in terms of runtime (Table 3, Table 4).

As the lookup embeddings or HMM profiles are computed only once, we neglected this additional step. Such preparations cost much more than single queries: pre-computing HMM profiles using MMseqs2 took 15m, pre-computing embeddings about 23m (Table 3) using the same set and machine but utilizing CPUs in one (MMseqs2) and GPUs in the other (ProtT5). Only MMSeqs2 generated and indexed its database rapidly (19.5s). However, pre-processing is required only once, rapidly amortizing when running many queries. The ability to pre-compute such representations is also a crucial difference between ProtTucker and other learned methods (16,68). For pairwise protein comparisons, those methods typically require N comparisons/forward-passes to search with a single query against N proteins. Instead, ProtTucker only needs a single forward pass to embed the new query; subsequent similarity scoring simply and quickly computes an Euclidean distance.

This makes ProtTucker search speed scale well with database growth suggesting the tool as a fast but sensitive pre-filter for other methods that in turn provide residue-level information as showcased on three model organisms (Table 4), including one of the largest organisms on earth (fungus *A. Ostoyae*, 22,192 proteins) and one of the largest viruses (*M. Chilensis*, 1,120 proteins). In less than 27 min on a single machine (Table 4), ProtTucker transferred substantially more CATH annotations mapping proteins from their sequence to 3D structures through the CATH resource than Gene3D (5) at a similar level of expected error (Table 4).

For the virus and the bacterium (*E. coli*) fungus the annotations agreed to over 80% with Gene3D, while this value dropped to 75% for the fungus (Table 4). Although high, the agreement was lower than expected: if ProtTucker and Gene3D each had fewer than 5% errors, then both should agree for over 90% of the proteins for which both transfer annotations. Most likely, this discrepancy (Δ(90-80)) arose partially from multi-domain proteins. Despite carefully cross-validating ProtTucker, an alternative explanation for the discrepancy is underestimating the expected error a distances ≤0.9 as 5% instead of up to 15%. The “functional shape” of the agreement between ProtTucker and Gene3D at different distance thresholds (Table S7) suggested that the “errors” (lack of agreement) did not only originate from ProtTucker. Carefully annotating the five proteins with the lowest distance and a different CATH annotation (Table S4) supported this perspective.

The agreement for multi-domain proteins dropped less than expected (11 percentage points drop for *M. Chilensis*, 5 percentage points increase for *E. coli*), possibly suggesting that ProtTucker using averages over an entire protein for comparison did not trip substantially more over the multi-domain challenge than the local alignment-based Gene3D using HMMER (6). This might suggest ProtTucker to have added correct annotations over Gene3D in multi-domain proteins, although developed exclusively on single domain proteins. The substantial increase in coverage from the level expected at distances ≥0.9 (Fig. 5, Table S4) for the proteomes (Table 4) might be misleading: to establish performance coverage (Fig. 5, Table S4), we used a highly non-redundant lookup set, presumably removing many easy hits. In contrast, analyzing proteomes, we transferred annotations for all CATH-S100 proteins, leveraging “redundant annotation transfers” to increase coverage.

As for HBI, the accuracy of EAT also increased for larger families (Fig. S1). One explanation is that the larger the family, the higher the random hit rate, simply because there are more possible hits. Another, more subtle (and given the enormous compute time needed to train ProtT5, more difficult to test) explanation is that the largest CATH families represent most of the largest protein families (2). In fact, a few hundred of the largest superfamilies cover half of the entire sequence space (2,88). Simply due to their immense size, these large families have been sampled more during the pre-training of ProtT5.

### ProtTucker embeddings intruded into midnight zone

The embedding space resulting from contrastive learning, introduced here, improved performance consistently for all four pLMs (Table 1). This was revealed through several ways of looking at the results from embeddings with and without contrastive learning: (1) the increased separation of protein pairs within the same protein superfamily and between different superfamilies (Fig. 2), (2) the qualitative improvement in the clustering of t-SNE projections (Fig. 3), the better correlation of embedding distance and structural similarity (Fig. 4) and (3) the quantitative improvement in the EAT benchmark (Table 1). On top, the Euclidean distance correlated with accuracy (Fig. 5, Table S3). Similar to an E-value in HBI, this lets users gauge the reliability of a hit between query and annotated protein.

While the accuracy of the best performing pLM (ProtTucker(ProtT5)) was similar to HBI using HMM-profiles on the most fine-grained level of homologous superfamilies (CATH level H, Table 1), the relative advantage of EAT increased, the more diverged the level of inference, i.e., EAT outperformed HBI for more distant relations from the midnight zone (CATH level C, Table 1). When further reducing data redundancy, i.e., removing more similar sequences, this trend became clearer (Fig. 6). Despite increasing difficulty, the performance of EAT decreased almost insignificantly where HBI approached random for insignificant E-values. This trend was supported by the correlation of structural similarity as defined by SSAP (7,8) and the Euclidean distance between protein pairs in a 30% non-redundant data set (Fig. 4).

ProtTucker and tools such as HMMer have very different resolution: ProtTucker considers only per-protein averages to match query to template. In contrast, HMMer - or similar methods - align each residue between both proteins. The coarse-grain yields the speedup (Table 4), and pitches ProtTucker as a fast pre-filter. Once the hit is found by scanning large data sets, the slower, fine-grained methods for per-residue alignments and 3D prediction can be employed. However, the per-protein average also implies limitations, e.g., when Q and T have very different numbers of domains or the number of domains for Q is not known (Table 3).

Ultimately, the coarse-grained ProtTucker can compete at all because embeddings intrinsically abstract the constraints under which protein sequences evolve, including constraints upon structure, function, and the environment. The same constraints coin the evolutionary information contained in profiles of protein families. Apparently, pLMs such as ESM-1b (12), ProtBERT (18), or ProtT5 (18) are successfully condense these constraints. In fact, pLMs are arguably more successful than profile-based methods because a simple length-average over the position-specific scoring metrices (PSSM) would not suffice to predict CATH numbers very accurately.

ProtTucker builds upon this success to explicitly capture the constraints relevant for the CATH hierarchy. Thus, the less a particular aspect of function depends on structure, the less likely the new ProtTucker embeddings will reflect this aspect. On the other hand, an approach similar to ProtTucker focused on particular functional hierarchies, e.g., EC numbers appears to work well (SM Akmese & M Heinzinger, unpublished).

Taken together, these results indicated that contrastive learning captured structural hierarchies and provides a novel, powerful tool to uncover structural similarities clearly beyond what has been achievable with 50 years of optimizing sequence-based alignment techniques. Using EAT to complement HBI could become crucial for a variety of applications, ranging from finding remote structural templates for protein 3D structure predictions over prioritizing new proteins without any similarity to an existing structure to filtering potentially wrong annotations. One particular example has recently been shown for the proteome of SARS-CoV-2 to unravel entire functional components possibly relevant for fighting COVID-19 (70).

### ProtTucker embeddings improved FunFams clustering

Previously (59), we showed that a simplistic predecessor of ProtTucker helped to refine the clustering of FunFams (79). By adding an additional, more fine-grained hierarchy level in CATH, FunFams link the structure-function continuum of proteins. The functional consistency within FunFams was proxied through the enzymatic activity as defined by the EC (Enzyme Commission (82)) number. Even the preliminary ProtTucker improved the annotation transfer of ligand binding and EC numbers (59) by removing outliers from existing FunFams and by creating new, more functionally coherent FunFams. As for CATH, the contrastively trained ProtTucker(ProtBERT) also improved over its unsupervised counterpart, ProtBERT, for FunFams. It improved functional consistency especially for proteins in the twilight zone (<35% PIDE, Fig. 5 in (59)). Thus, ProtTucker embeddings improved functional (FunFams) and structural (CATH) consistency beyond sequence similarity. Here, we expanded upon this analysis by showing how EAT can be improved even more through contrastively learning hierarchies. Using the proposed method, we could spot potential outliers, i.e., samples with the same annotation but large embedding distance. This might become essential to clean up databases. Aside from outlier-spotting, we could also obtain labels from the nearest neighbors of outliers (Table S6). Although we could not reproduce the same level of success when applying EAT to inferring subcellular location in ten states (Table 3), the CATH-optimized ProtTucker embeddings also did not perform worse.

### Generic advantages of *contrastive learning*

Contrastive learning benefits from hierarchies as opposed to supervised training which usually flattens the hierarchy thereby losing its intrinsic advantage. Other possible advantages of contrastive learning include the following three. (1) <u>Dynamic data update (*online learning*):</u> While supervised networks require re-training to benefit from new data, contrastively trained networks can benefit from new data by simply updating the lookup set. This could even add completely new classes, such as proteins for which the classification will become available only in the future. HBI shares this advantage that originates from the difference between classifying proteins into existing families versus classifying by identifying the most similar proteins in that family. (2) <u>Learn the access, not the data:</u> Instead of forcing the supervised network to memorize the training data, contrastive learning teaches how to access the data stored in an external lookup set. (3) <u>Compression:</u> As many other learning techniques, contrastive learning can act as a compression technique. For instance, we reduced the disk space required to store protein embeddings threefold by projecting 1024-dimensional vectors onto 128 dimensions while improving performance (Table 1). This renders new queries (inference) more efficient and enables scaling up to very large lookup sets. (4) <u>Interpretability:</u> knowing from which protein an annotation was transferred might help users benefit more from a certain prediction than just the prediction itself. For instance, knowing that an unnamed query protein shares all CATH levels with a particular glucocorticoid receptor might suggest some functional implications helping to design future experiments.

Gaining from embeddings for sequence comparisons. The solution described here established the power for contrastive learning. Other, more straightforward solutions

## Conclusions

Embeddings from protein Language Models (pLMs) extract the information learned by these models from unlabeled protein sequences. Embedding-based Annotation Transfer (EAT) replacing the proximity in sequence space used by homolog-based inference (HBI) through proximity in embedding space already reaches traditional alignment methods in transferring CATH annotations from a template protein with experimental annotations to an unlabeled query protein. Although not quite reaching the performance of advanced profile-profile searches by HMMer for all four CATH levels, the best embeddings surpassed HMMer for two of the four levels (C and A). When optimizing embeddings through contrastive learning for the goal of transferring CATH annotations, EAT using these new embeddings consistently outperformed all sequence comparison techniques tested. This higher performance was reached at a fraction (three orders of magnitude) of the computational time. Although the new embeddings optimized through contrastive learning for CATH did not improve performance for a completely different task, namely the prediction of subcellular location in ten classes, the CATH-optimized solution did also not perform significantly worse. Remarkably, just like HBI, the performance of EAT using the optimized ProtTucker embeddings was proportional to family size with increased accuracy for larger families.

## Supporting information

Supplementary Online Material (SOM)

## Availability

Building on top of bio_embeddings package (89) we have made a script available that simplifies EAT https://github.com/Rostlab/EAT.

## Abbreviations used

3D: three-dimensional
BFD: Big Fantastic Database (11)
CATH: hierarchical classification of protein 3D structures in Class, Architecture, Topology and Homologous superfamily (1,2)
DL: Deep Learning
EAT: Embedding-based Annotation Transfer
EI: evolutionary information
embeddings: fixed-size vectors derived from pre-trained pLMs
ESM-1b: pLM from Facebook dubbed Evolutionary Scale Modeling (12)
FNN: Feed-forward Neural Network
FunFams: functional families as sub-classification of the most fine-grained H level in CATH (13)
HBI: Homology Based Inference
HMM: Hidden Markov Model
HMMer: particular method for HMM-profile alignments (6)
HSSP: homology-derived secondary structure of proteins (14)
HVAL: distance from empirical curve separating proteins with similar structure recognizable from pairwise alignments (15)
LM: Language Model
MMseqs2: fast database search and multiple sequence alignment method (10)
MSA: Multiple Sequence Alignment
NLP: Natural Language Processing
PDB: Protein Data Bank
PIDE: percentage pairwise sequence identity
pLM: protein Language Model
ProSE: pLM based on long short-term memory (LSTM) cells dubbed Protein Sequence Embeddings (16)
ProtBERT: pLM based on the LM BERT (17)
ProtT5: pLM based on the LM T5 (18)

## Acknowledgements

Thanks primarily to Tim Karl, but also to Guy Yachdav & Laszlo Kajan (all TUM) for invaluable help with hardware and software; to Inga Weise (TUM) for support with many other aspects of this work. Many thanks for both anonymous reviewers, deep kudos for the detailed help from No. 1! This work was supported by the Bavarian Ministry of Education through funding to the TUM and by a grant from the Alexander von Humboldt foundation through the German Ministry for Research and Education (BMBF: Bundesministerium für Bildung und Forschung), by two grants from BMBF (031L0168 and program “Software Campus 2.0 (TUM) 2.0” 01IS17049) as well as by a grant from Deutsche Forschungsgemeinschaft (DFG-GZ: RO1320/4-1). Last, but not least, thanks to all those who maintain public databases in particular Steven Burley (PDB, Rutgers), Alan Bridge (Swiss-Prot, SIB, Lausanne), Alex Bateman (UniProt, EBI Hinxton) and their crews, and to all experimentalists who enabled this analysis by making their data publicly available.

