## Supplementary Online Material (SOM) for "Contrastive learning on protein embeddings enlightens midnight zone"

#### Table of Contents for Supporting Online Material

#### Short description of Supporting Online Material

In this Supporting Online Material (SOM), we provide more information on the effect of family size on accuracy when using embedding-based annotation transfer (EAT) to predict homologous superfamilies in CATH (*1, 2*) (Fig. S1). Additionally, we show how the number of samples in our validation (val200) and test set (test219) changes with respect to the hierarchy level in CATH (Table S1). Along the same line, we add details on the number of unique CATH classes at the four levels of the hierarchy for train66k, val200 and test219 (Table S2). We also provide more details on Fig. 5 allowing users to make an informed decision about the choice of the Euclidean distance with respect to expected accuracy (Table S3) and coverage (Table S4). In order to probe to which extent the newly learnt embeddings generalize to other problems, we also compared EAT performance for the task of subcellular localization prediction before/after applying contrastive learning (Table S5). Based on our finding that the Euclidean distance between embeddings gives information on the reliability of an EAT prediction (Fig. 5), we also provide a use-case that exemplarily demonstrates how embedding distance can be useful to detect outliers in current annotations (Table S6). Lastly, we investigate the Euclidean distance threshold at a proteome-scale by comparing the agreement between EAT and HBI predictions for three different proteomes at varying distance cut-offs.

### Material

**Fig. S1: Large families easier to predict:**

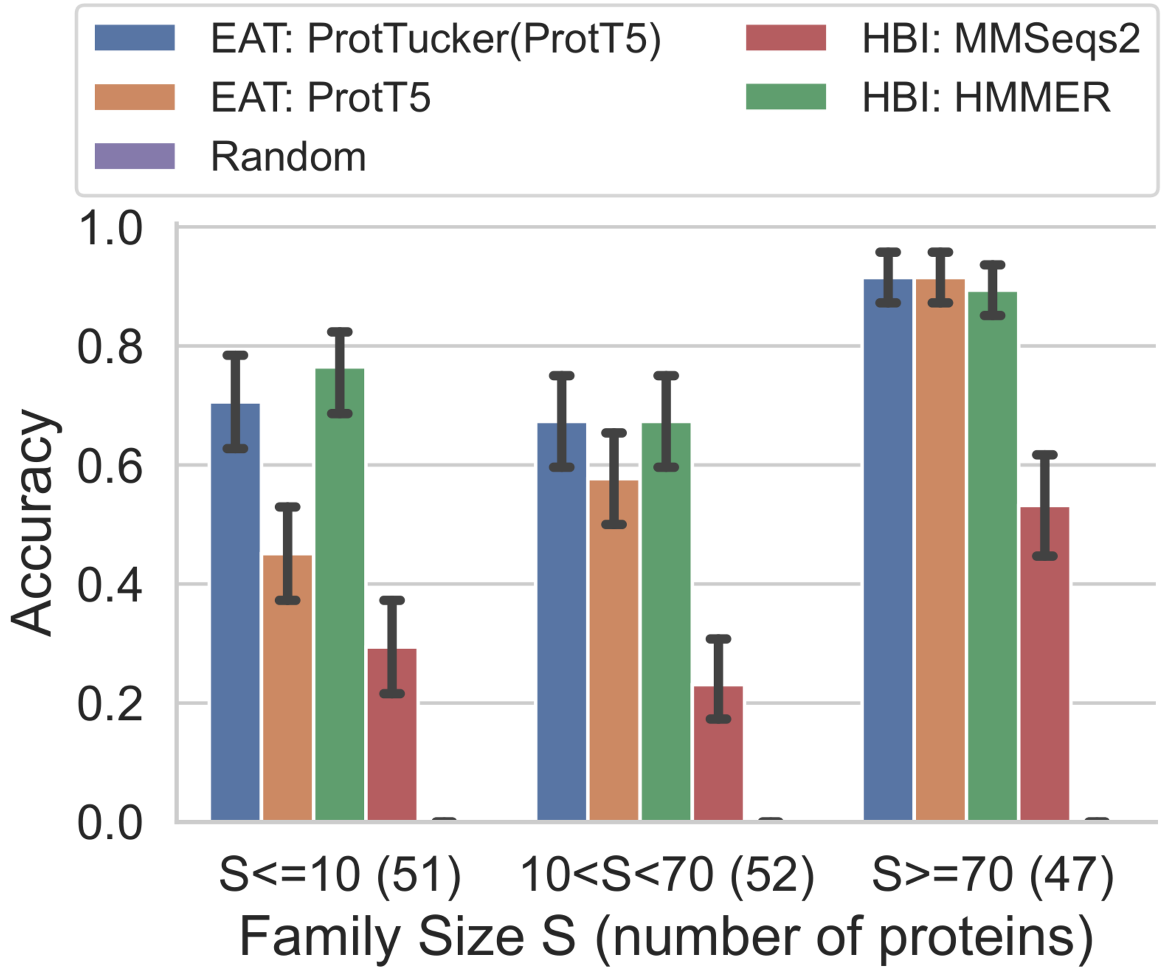

**Fig. S1: Large families easier to predict.** To analyze the effect of protein family size on performance, we transferred annotations from lookup69k to test219 and compared the fraction of correctly predicted homologous superfamilies (accuracy, Eqn. 3) between three groups of family sizes S: small families (S≤10 in lookup69k; 51 proteins), medium-sized (10<S<70; 52 proteins) and large (S≥70; 47 proteins). We compared two HBI-based methods (red: MMSeqs2 (*3*) and green: HMMER (*4*)) and two EAT-based methods (orange: raw ProtT5 (*5*) and blue: new ProtTucker(ProtT5)). If a method did not provide any prediction, this sample was counted as incorrectly classified. Error bars indicated 95% confidence intervals computed via bootstrapping (Eqn. 6). Sequence-based MMSeqs2 performed significantly worse than the other methods because it was used for redundancy reduction. The optimized ProtTucker(ProtT5) embeddings introduced here, improved over the unsupervised ProtT5, especially, for small-sized families, reaching performances on par with HMM-based HMMer without requiring MSAs as input.

**Table S1: Number of proteins in data sets by CATH-levels ***

| | C | A | T | H |
|---|---|---|---|---|
| val200 | 200 | 200 | 190 | 153 |
| test219 | 219 | 219 | 210 | 150 |

* Only a subset of the proteins in our final test-set (test219) or our validation set (val200) had at least one other protein in the Lookup69k or train66k set with the same CATH annotation at a certain CATH hierarchy level. We excluded test219/val200 proteins without a partner from the evaluation at a certain CATH level because they could not be predicted correctly. This table shows how the test-set size varies for the different CATH-levels.

**Table S2: Number of classes in data sets by CATH-levels ***

| | C | A | T | H |
|---|---|---|---|---|
| train66k | 5 | 43 | 1446 | 6479 |
| val200 | 4 | 23 | 125 | 153 |
| test219 | 5 | 22 | 123 | 150 |

* Only a subset of CATH classes was present in our final test-set (test219) and validation set (val200). Further, we excluded test219/val200 proteins that did not have had at least one other protein in the Lookup69k/train66k set with the same CATH annotation at a certain CATH hierarchy level because they could not be predicted correctly. This table shows how the number of classes varies for the different CATH-levels.

**Table S3: Accuracy depending on Euclidean distance threshold***

| Threshold | C | A | T | H |
|---|---|---|---|---|
| 0.8 | 100 | 100 | 100 | 100 |
| 0.9 | 100 | 97 | 96 | 95 |
| 1.0 | 100 | 96 | 95 | 93 |
| 1.1 | 96 | 93 | 91 | 90 |
| 1.2 | 96 | 89 | 84 | 87 |

| | | | | |
|---|---|---|---|---|
| 1.3 | 92 | 84 | 74 | 85 |
| 1.4 | 89 | 79 | 67 | 83 |

* Similar to varying E-value cut-offs for HBI, we examined whether the fraction of correct predictions (accuracy; Eqn. 3) depended on embedding distance for EAT. This was shown by transferring annotations for all four CATH levels from lookup69k to the queries in set test219 using the hit with lowest Euclidean distance. The fraction of test219 proteins having a hit below a certain distance threshold was evaluated separately for each CATH level. As decreasing embedding distance correlated with EAT performance, users can select an accepted error rate, e.g., 5% for hits below an Euclidean distance of 0.9.

**Table S4: Coverage depending on Euclidean distance threshold ***

| Threshold | C | A | T | H |
|---|---|---|---|---|
| 0.8 | 20 | 20 | 20 | 28 |
| 0.9 | 34 | 34 | 36 | 48 |
| 1.0 | 47 | 47 | 49 | 66 |
| 1.1 | 57 | 57 | 59 | 75 |
| 1.2 | 70 | 70 | 72 | 88 |
| 1.3 | 84 | 84 | 84 | 93 |
| 1.4 | 94 | 94 | 94 | 97 |

* Similar to varying E-value cut-offs for HBI, we examined whether the fraction of predictions depended on embedding distance for EAT. This was shown by transferring annotations for all four CATH levels from lookup69k to the queries in set test219 using the hit with lowest Euclidean distance. The fraction of test219 proteins having a hit below a certain distance threshold (coverage, Eqn. 4) was evaluated separately for each CATH level. As decreasing embedding distance correlated with EAT performance (SOM Table 3) and coverage, users can decide on the trade-off between coverage and performance based on their specific task.

**Table S5: EAT for subcellular location ***

| | 10-state subcellular location |
|---|---|

| | Accuracy (Q10) | Balanced accuracy | F1 |
|---|---|---|---|
| Random | 16 ±3 | 11 ±4 | 0.16 ±0.03 |
| ProtT5 | 54 ±4 | 43 ±6 | 0.54 ±0.05 |
| ProtTucker(ProtT5) | 52 ±2 | 42 ±6 | 0.53 ±0.04 |

* **Generalist vs Specialist:** Accuracy, balanced accuracy and F1 as implemented in SciKit (*6*) for predicting the subcellular localization of a protein in 10-states using embedding-based annotation transfer (EAT) via the pLM ProtT5 (generalist) or the proposed ProtTucker optimized on structural similarity (specialist). We used smallest Euclidean distance in embedding space as means to transfer annotations from proteins *DeepLocTrain (7)* to proteins in *setHard* (*8*). A random baseline was established by transferring the annotation of a random protein in *DeepLocTrain* to *setHard.* While the proposed ProtTucker outperforms the pLM ProtT5 on the task it was optimized for (structural similarity; Table 1), there appears to be neither a significant gain nor loss of performance when predicting other aspects of proteins such as showcased here using prediction of subcellular localization in 10-states.

**Table S6: Use-case: outlier detection** *

| | CATH ID-1 | CATH ID-2 | CATH ID-1/ID-2 | NN ID-1 | NN ID-2 | CATH NN ID-1 | CATH NN ID-2 |
|---|---|---|---|---|---|---|---|
| 1 | 1gkh A00 | 3kyh C02 | 2.40.50.140 | 1vqd A00 | 4pz7 A01 | 2.40.50.14 | 2.40.50.14 |
| 2 | 2zr1 A02 | 2g5x A02 | 4.10.470.10 | 2rde B01 | 1u0j A01 | 2.30.110.10 | 1.10.10.950 |
| 3 | 1cf7 A00 | 1ev7 B02 | 1.10.10.10 | 1cf7 B00 | 3v67 A01 | 1.10.10.10 | 3.30.450.210 |
| 4 | 4pdy A01 | 3skj F00 | 3.30.200.20 | 2q83 A01 | 3hei A00 | 3.30.200.20 | 2.60.120.260 |
| 5 | 4qva A02 | 3r6k A02 | 2.40.30.120 | 4qv2 A02 | 3ihm A02 | 2.40.30.120 | 3.30.9.40 |

* Outlier detection: One potential advantage of the correlation between embedding distance and prediction performance (Fig. 5) is that proteins that are annotated to be within the same superfamily but have a large embedding distance could point towards potential outliers in a protein family. Towards this end, we computed pairwise Euclidean distances between all *train66k* proteins within the same superfamily and manually checked the five pairs with the same superfamily but the largest distance (sorted in descending order in Table 4). Additionally, we

computed nearest neighbors (NN) for all samples to get candidates for alternative assignment. While the second (Ricin subunit) and last pair (Positive stranded ssRNA viruses) refer to relatively small families, the other pairs originate from large and diverse families responsible for Nucleic-acid binding (ID=1,3) or Kinases (ID=4). In the first example, one domain is annotated to be DNA-binding while the other is labelled as RNA-binding. These related yet different functions, could explain the large embedding distance despite having the same superfamily annotation.

Despite being annotated to the same superfamily, visual comparison of the second example suggests better structural agreement between 2g5xA02 and its NN (1u0jA01), compared to the structure of its superfamily-partner, 2zr1A02. The same holds true for the fourth example: while the protein pair is annotated to the same superfamily (Phosphorylase Kinase), the NN of 3skjF00 (3heiA00) appears structurally more similar to its NN. Additionally, both proteins, 3skjF00 and 3heiA00, are mapped to the same UniProt entry to transfer protein function as defined by their E.C. number (Enzyme Classification, (*9*)) suggesting some functional similarity that is detected by EAT but currently not reflected in CATH.

**Table S7: Threshold-dependency of proteome-analysis** *

| Δ | *E. Coli (K12)* | | | *A. Ostoayae* | | | *M. Chiliensis* | | |
|---|---|---|---|---|---|---|---|---|---|
| | Agree. [%] | Agree.-multi [%] | Cov. [%] | Agree. [%] | Agree.-multi [%] | Cov. [%] | Agree. [%] | Agree.-multi [%] | Cov. [%] |
| 1.1 | 81 | 86 | 100 | 70 | 57 | 98 | 82 | 73 | 98 |
| 1.0 | 81 | 86 | 99 | 72 | 60 | 92 | 83 | 73 | 96 |
| 0.9 | 81 | 86 | 97 | 75 | 65 | 79 | 84 | 73 | 87 |
| 0.8 | 81 | 87 | 91 | 81 | 74 | 51 | 88 | 77 | 60 |
| 0.7 | 83 | 88 | 80 | 89 | 87 | 27 | 94 | 94 | 29 |
| 0.6 | 86 | 90 | 68 | 93 | 92 | 17 | 91 | 86 | 11 |
| 0.5 | 90 | 94 | 50 | 96 | 95 | 9 | 96 | 100 | 4 |
| 0.4 | 94 | 95 | 34 | 98 | 90 | 4 | 86 | - | 1 |

* Threshold-dependency of proteome-analysis: Comparison of the annotation-transfer from 123k CATH-S100 proteins through HBI (Gene3D) and through EAT as introduced here (ProtTucker(ProtT5), or PT(ProtT5)) for three entire reference proteomes: Escherichia coli (E. Coli), Armillaria ostoyae (A. ostoyae) and Megavirus Chilensis (M. Chilensis). In other words, all proteins in the three organisms were mapped to proteins of known structure using the CATH hierarchy. Gene3D predictions were taken from UniProt; PT(ProtT5) predictions were derived from the single nearest neighbor in Euclidean space up to a maximum distance threshold Δ ranging from 1.1 to 0.4. Agreement (Agree.): fraction of proteins for which Gene3D

and PT(ProT5) had a prediction and reported the same homologous CATH superfamily (for multi-domain proteins with multiple Gene3D annotations, matching any domain by PT(ProtT5) was considered as correct); Agreement multi-domain (Agree.-multi): fraction of multi-domain proteins for which the homologous CATH superfamily predicted by PT5 agreed with one of the Gene3D domain annotations; Coverage (Cov.): the fraction of proteins in the proteome for which EAT provided predictions with a distance less than Δ. At very low distance thresholds, i.e., below 0.6, the steep decrease in coverage can lead to reduced agreement compared to larger distance thresholds as the number of predictions for which EAT and HBI provided predictions decreased.
